# Spatial Attention and Session Day Independently Modulate Human Visual Cortical Plasticity

**DOI:** 10.1101/2025.10.31.685945

**Authors:** Patricia Naomi Limon, Anthony M. Norcia, Ryan Thomas Ash

## Abstract

Stimulus-specific response potentiation (SSRP) is a noninvasive form of cortical neuroplasticity elicited by repeated presentation of high-contrast visual stimuli. Analogous to long-term potentiation, SSRP has been proposed to exhibit input specificity, with potentiation confined to neural populations driven by the induction stimulus. In rodent models, SSRP effects accumulate across days; however, it remains unclear whether selective attention influences the magnitude or specificity of potentiation in humans. Here, we examined whether covert spatial attention modulates SSRP strength using high-density electroencephalography (EEG) in neurotypical adults. An established SSRP paradigm was modified to include an attention task during induction. Pre- and post-induction amplitudes were measured using frequency-tagged (6 and 7.5 reversals/s) bilateral hemifield contrast-sweep checkerboard steady-state visual evoked potentials (SSVEPs). The 10-minute, high-contrast, 2 Hz sign-reversing induction stimulus was presented exclusively to the left hemifield, with the right serving as control. Across two experimental sessions, participants attended either toward or away from the potentiated hemifield during induction. SSRP produced increased post-induction amplitudes in both hemifields, challenging the notion of strict input specificity. Potentiation was significantly greater on the second session day, independent of attention condition. Notably, attention enhanced SSRP in naïve but not experienced observers, reflecting a significant interaction between attention and session day. Together, these findings suggest that (1) human SSRP may not be strictly stimulus-specific, (2) attention modestly enhances SSRP during initial exposure, and (3) repeated induction produces a robust plasticity effect that occludes attentional modulation.

## Introduction

One established approach to non-invasively probe visual cortical plasticity in humans and other species is through stimulus-specific response potentiation (SSRP), characterized by enhanced visual evoked potentials (VEPs) following a period of repetitive exposure to a stimulus. It has been demonstrated that a stimulus-driven plasticity response of visual evoked potentials can be induced through passive viewing of a high-contrast sign-reversing stimulus for as little as 2-10 minutes (Teyler et al., 2005, Frenkel et al., 2006, Normann et al., 2007, Kirk et al., 2008, Cooke and Bear, 2010, Valstad et al., 2021). Its induction through sensory experiences makes SSRP a valuable model for studying functional brain plasticity in vivo. SSRP mechanisms may contribute to facilitating certain forms of perceptual learning which, broadly, involves the organization, identification, and interpretation of sensory information due to visual exposure (Lu et al., 2011, Donovan et al., 2020, Hung and Carrasco, 2021). Through these processes, the brain develops a heightened ability (lowered threshold) to detect target stimuli, potentially utilizing synaptic mechanisms similar to those observed in long-term potentiation (LTP). SSRP shares several characteristics with Hebbian LTP, particularly dependency on NMDA receptor activation and input specificity (Kirk et al., 2008, Cooke and Bear, 2010, Kirk et al., 2021). Treatments that extinguish LTP, such as local expression of GluR1-CT peptide or administration of zeta inhibitory peptide, also disrupt SSRP, suggesting that SSRP depends on this form of synaptic plasticity and that its modulation may provide an in vivo measure of LTP-like processes in the human cortex (Frenkel et al., 2006, Cooke and Bear, 2010). Since SSRP and LTP mechanisms resemble each other in terms of synaptic transmission development, quantifying SSRP susceptibility holds the potential to serve as a biomarker in neuropsychiatric illness. For example, individuals with bipolar disorder (Type I and II), major depressive disorder, and schizophrenia have been shown to exhibit impairments in SSRP (Normann et al., 2007, Jacob et al., 2021b, Valstad et al., 2021). These findings suggest that noninvasive LTP-like plasticity probes, such as SSRP, could serve as biomarkers in clinical trials for these illnesses.

In rodents, SSRP-induced response enhancement is highly selective, as slight variations in the spatial frequency, orientation, or visual contrast of the experienced stimulus do not elicit the same potentiated response, highlighting its input specificity (Frenkel et al., 2006, Kirk et al., 2008, Cooke and Bear, 2010). Additionally, SSRP induced in one eye does not elicit the same VEP response potentiation in the untrained eye (Frenkel et al., 2006), suggesting that it arises at early stages of the visual pathway, prior to binocular integration. However, these findings are based on non-human studies, raising questions about their generalization to humans, given species-specific differences in visual system organization. To our knowledge, ocular transference of SSRP has not been investigated in humans, and while SSRP has been shown to be selective for orientation, contrast, and eye of origin, its retinotopic specificity, a logical extension of these properties, remains under-explored (but see Teyler et al., 2005). Here, we examine input specificity by inducing SSRP in one visual hemifield and measuring visual evoked responses to stimuli in both hemifields, before and after SSRP induction. Since SSRP reflects experience-dependent plasticity in visual cortex, it is important to consider how top-down factors such as attention may modulate its expression. Interestingly, SSRP paradigms have yet to be examined alongside established modulators of perceptual learning and learning more broadly. Attentional state plays a fundamental role in shaping learning and memory, particularly perceptual memories (Mukai, 2011, Szpiro and Carrasco, 2015, Donovan and Carrasco, 2018). In perceptual learning paradigms, attention has been shown to enhance neural and behavioral responses to target stimuli (Nguyen et al., 2020, Cavanaugh et al., 2022). Moreover, paired associative stimulation (PAS)-induced plasticity in human motor cortex is enhanced when the stimulated limb is attended visually and spatially, compared to when attention is allocated elsewhere (Stefan et al., 2004). Follow-up studies of PAS show that attention exhibits opposite effects depending on induction protocol: it enhances motor evoked potentials (MEPs) during LTP-like stimulation but reduces MEP depotentiation during LTD-like stimulation, indicating bidirectional modulation of neuroplasticity (Kamke et al., 2012). These findings suggest that attention may act as a general modulator of experience-dependent synaptic change across cortical systems, including SSRP. To investigate this in the visual system, we examined covert endogenous spatial attention (a form of voluntary, stimulus-independent orienting toward specific retinotopic locations) (Carrasco, 2011) to determine its influence on visual cortical plasticity.

Another important feature of SSRP, observed thus far in rodents, is that potentiation accumulates across experimental days (Frenkel et al., 2006, Cooke and Bear, 2010), suggesting that consolidation processes involving sleep and time contribute to recalibrating neural circuitry strength. In humans, however, previous reports describe potentiation effects emerging within minutes after a visual plasticity protocol (Teyler et al., 2005, Normann et al., 2007, Klöppel et al., 2015, Valstad et al., 2020). Our laboratory has not yet observed such immediate effects, as our initial SSRP induction study did not yield potentiation (sign-reversing gratings at 18 reversals/second presented in contrast sweeps for 900 seconds) (Ash et al., 2023), highlighting a broader issue with the paradigm: its replicability across laboratories remains inconsistent, possibly due to variability in participant population or details of visual stimulus presentation (Çavuş et al., 2012, Valstad et al., 2020, Sumner et al., 2020, Jacob et al., 2021a, Dias et al., 2022). Here we employed steady-state visually evoked potentials (ssVEPs) due to its high signal-to-noise readout of population activity and its unique ability to enable simultaneous recording activity in both hemifields with frequency-tagging (Morgan et al., 1996, Müller et al., 1998, Norcia et al., 2015, Ash et al., 2023). These characteristics of the ssVEP enable in-vivo quantification of how attention shapes experience-dependent plasticity.

### Hypotheses

Based on prior studies in rodents (Frenkel et al., 2006, Cooke and Bear, 2010) and humans (Teyler et al., 2005), we predict that (I) SSRP will produce response potentiation in the potentiated but not control hemifield, reflecting LTP-like retinotopic specificity. Building on the results of previous studies (Stefan et al., 2004, Ling and Carrasco, 2006, Donovan et al., 2020, Hung and Carrasco, 2021), we further predict that (II) attentional deployment toward the induction stimulus (attention–induction congruent condition) will lead to greater plasticity—indexed by increased VEP amplitudes— than when attention is deployed away from (contralateral to) the induction stimulus (attention–induction incongruent condition). Finally, based on Cooke and Bear and Frenkel’s SSRP investigations, we predict that (III) response amplitudes in the potentiated hemifield will increase across repeated days of SSRP, consistent with cumulative effects over time.

## Methods

### Participants

Thirty-seven participants were recruited from Stanford University’s research pool databases. Individuals 18 years of age or older, with no known neurological impairments and normal or corrected-to-normal vision, were eligible to participate. Informed consent was provided for all participants prior to any data collection. The study was approved by the Stanford IRB. Participants completed two EEG sessions and received course credit (1 credit / 1 hour).

### Exclusion Criteria

EEG session data was excluded if more than 30% of signal data quality was above participant signal threshold across all contrasts (n = 11). A total of twenty-six participant’s EEG data was analyzed.

### Data Acquisition

Prior to arrival, participants were instructed to wash their hair with shampoo (no conditioner). Vision testing was performed to confirm normal or corrected-to-normal vision. Afterwards, high-density, 128-channel electroencephalograms (EEG) were recorded using HydroCell electrode arrays and an Electrical Geodesics Net Amps 400 (Electrical Geodesics) amplifier. The EEG cap was prepared in a KCl solution for 10 minutes. Electrode caps were placed on the participant’s head, and additional KCl solution was applied to individual electrodes such that all electrodes had an impedance < 50 kOhm. Impedance checks were performed twice during the recording session, each session lasting 45 minutes.

### Visual Stimuli and Attentional Paradigm

Visual stimuli were displayed on a high dynamic range 60-Hz refresh rate monitor positioned 70 cm from the participant’s eye level. Stimuli were generated with custom lab software XDiva. Stimuli consisted of two semicircular checkerboards each positioned laterally on each side of fixation with the stimuli spanning 0.5° to 10° eccentricity. Each checkerboard wedge was divided into sixteen 0.628° segments along the vertical meridian relative to fixation, and 20 rings with a spatial frequency of 0.5°, creating a wedged checkerboard pattern. The semicircular design left the central area near fixation devoid of pattern elements, and both this region and the background were maintained at mean background luminance. Each hemifield stimulus was frequency-tagged to measure EEG ssVEPs. Pre-/post-stimuli sign-reversal rates were F1 = 3 Hz (6 reversals-per-second, rps, left hemifield), F2 = 3.75 Hz (7.5 rps, right hemifield). Pre-/post-stimuli were contrast sweep stimuli presented in log steps of: 1%, 2.5%, 6.3%, 15.8%, 39.8% and 100%. Each contrast level was presented for 1.33 s (8 s of core stimulus bins). Each stimulus sweep began with a 1.33 s prelude bin at 1% contrast and ended with a 1.33 s postlude bin at 100% contrast. These stabilization periods help reduce edge artifacts in the broadband EEG for subsequent Fourier analysis. The total stimulus duration, including prelude/postlude was 10.66 s. There was a 3000 ms ± 500 ms inter-trial rest period between each stimulus. Participants were advised to limit blinking to inter-trial periods (confirmed by visual inspection of the EEG). For each session, 34-40 (generally 39) trials of contrast-sweep ssVEPs were acquired before SSRP, and another 34-40 trials were acquired after SSRP. The SSRP induction stimulus was modified from the pre-post-stimulus such that the left checkerboard was presented at 100% contrast (not swept) flickering at F1 = 1 Hz (2 rps) a parameter previously reported to induce neuroplasticity. The right checkerboard, serving as the non-potentiating control stimulus was presented at 2% contrast with frequency F2 = 1.5 Hz (3 rps). The stimuli were presented simultaneously for 30 seconds with a brief (<0.5 sec) gap, for 10 trials. The participant was then given a few seconds of rest, and the participant observed another ten 30-second trials, for a total SSRP induction duration of 10 minutes, in line with Norman et al.’s induction time (Normann et al., 2007). During the SSRP block, participants were advised to blink when they needed to but to not close their eyes for extended periods of time. A trial example for each condition can be observed in **Supplementary <u>Movies</u>**. In each session, participants completed a pre-SSRP Block, SSRP Block, and a post-SSRP block (**Fig. 1A**). During the pre-post-SSRP blocks, participants performed a simple letter detection task presented at the center of the screen to maintain fixation. A series of the letters L, F and T (0.25° retinotopic extent) were presented one at a time at the center of the screen approximately once per second at different orientations. Participants were instructed to make a response via a keyboard button when the letter T was presented, regardless of the orientation of the T. Attentional deployment was manipulated during the SSRP block (**Fig. 1B**). In the Potentiation-Attention Congruent condition, participants were instructed to fixate on the center of the screen and deploy peripheral spatial attention to the left checkerboard (SSRP stimulus) while 2°-square 100-ms contrast decrement patches appeared randomly within the left hemifield stimulus every 5-8 seconds. Participants were instructed to make a response by pressing a button on a keypad when a contrast decrement was detected. The contrast decrement started at 18%. Task difficulty increased following trials in which responses were made to all decrements, otherwise, task difficulty decreased, to maintain task difficulty at 70% percent correct. In the Potentiation-Attention Incongruent condition, participants were instructed to fixate on the center of the screen and deploy peripheral spatial attention to the right low-contrast checkerboard (non-induction stimulus) while 2°-square contrast increments appeared randomly within the right hemifield stimulus with otherwise similar features as in the attention congruent condition. The task increment started at 2% updated by staircase as above. Participants completed the two attention conditions on two separate days spaced 1-28 days apart (one participant completed the study after 38 days), the order of conditions was counterbalanced across participants.

**Figure 1.**
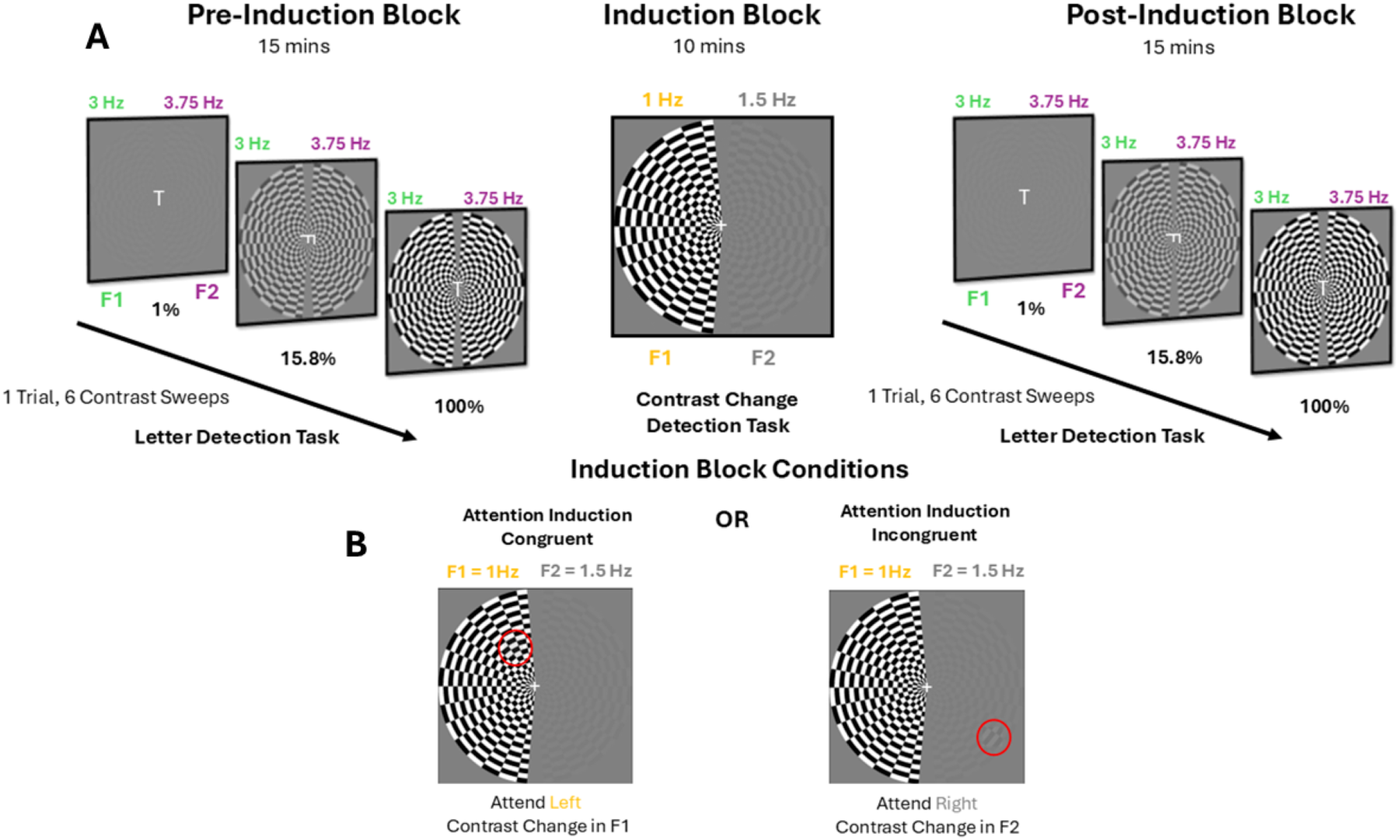
| Experimental Protocol. **A** Participants first completed a pre-SSRP block, 34-40 contrast-sweep ssVEP trials. Next, participants completed a 10-minute SSRP block. Here the left checkerboard served as the potentiating stimuli at 100% contrast sign-reversing twice a second and the right checkerboard served as the non-potentiating control (2% contrast, sign reversing at 3 rps). Finally, participants completed the post-SSRP block, which was identical to the pre-SSRP block. **B** Illustration of the two attentional deployment conditions during induction. Left panel: Attention induction congruent in which the participant attends ***toward*** the induction (left) stimulus. Right panel: Attention induction incongruent in which the participant attends ***away*** from the induction stimulus and toward the non-potentiating (right) stimulus, performing a contrast change detection task across 10 minute induction block. Attention conditions were performed in a counterbalanced order across experiment days.

### Signal Processing

The EEG was sampled natively at 500 Hz and then resampled at 420 Hz. Raw EEG data was processed with a 0.3-50 Hz bandpass filter with EGI system software. The data was next exported to the custom signal processing software XDiva. This software provided a digital trigger indicating the start of the trial with sub-millisecond accuracy. Artifact rejection was performed in two steps. First, the continuous filtered data were evaluated according to a sample-by-sample thresholding procedure to locate consistently noisy sensors. These channels were replaced by the average of their six nearest spatial EEG channels. Once noisy channels were interpolated in this fashion, the EEG was re-referenced from the Cz reference used during the recording to the common average of all sensors. Second, 1-s EEG epochs that contained a large percentage of data samples exceeding the threshold (30–80 microvolts) were excluded on a sensor-by-sensor basis. Afterwards, baseline correction 1333 ms prior to trial onset was performed to further increase the signal-to-noise ratio and reduce the effect of drifting. Finally, ssVEP amplitude and phase were calculated at the 2nd, 4th, 6th and 8th harmonic by a recursive least squares adaptive filter (Tang and Norcia, 1995), with weights for each component of complex values for each harmonic frequency analyzed (2F - 8F). To reduce the 128-channel data into interpretable, physiologically plausible linear components, we applied Reliable Components Analysis (RCA) adapted for SSVEPs. Trial-wise complex Fourier coefficients at the selected harmonics were computed for each condition. RCA seeks spatial filters that maximize trial-to-trial covariance (signal reliability) relative to within-trial covariance, formulated as a generalized eigenvalue problem. The resulting eigenvectors (spatial filters) are ordered by the amount of reliability they explain, and their forward models yield physiologically interpretable scalp topographies (Dmochowski et al., 2015). RCA weights were trained on a group-level concatenated multi-subject sensor-by-feature matrices and then projected back to each participant’s data individually through these weights to obtain component amplitudes/phases at each harmonic and contrast. This procedure emphasizes channels that contribute responses with consistent phase-locking across trials, increasing SNR and yielding components that closely match expected visual-cortical topographies. RCA was performed for each hemifield frequency tag independently, filtering at 2F1-8F1 harmonics (6, 12, 18, 24 Hz) and 2F2-8F2 harmonics (7.5, 15, 22.5, 30 Hz) as two separate runs. Depending on the hemisphere of interest, components were learned on RLS-filtered complex value data from either F1 or F2 responses across all sessions recorded (i.e. the same weighted sum of electrodes was used both pre and post SSRP measures). We note that for each hemifield frequency tag in our paradigm, the first RCA component falsely localizes to the ipsilateral occipital electrodes (**Supp. Fig. 1A**), likely due to dipole cancellation from the inferior and superior occipital sulcus (Ales et al., 2010). The first reliable component (RC) for each hemifield frequency tag was analyzed.

### Contrast Response Functions

For each participant, trials obtained before and after SSRP were averaged separately. Response amplitudes were then computed as the vector average of complex Fourier coefficients for each contrast and condition at each harmonic, yielding pre- and post-SSRP contrast response functions (CRFs). CRFs were derived for the 2nd, 4th, 6th, and 8th harmonics—the 2nd harmonic containing the primary response due to frequency doubling of the sign-reversing visual stimulus—separately for each session and hemifield. Next, to capture the total steady-state response across harmonics, root-mean-square (RMS) amplitudes were calculated, yielding a phase-independent measure of overall response magnitude. This procedure integrates the contribution of multiple harmonics, providing a stable estimate of neural response strength suitable for comparison across contrasts and experimental conditions. Finally, to enable pooling across participants, post-SSRP CRFs were normalized to the maximum pre-SSRP response amplitude within the same session and hemifield. This procedure scaled all post-induction responses relative to each participant’s own peak baseline response, controlling for variability in overall amplitude across sessions.

### Statistical analysis for Linear Mixed-Effects Model

Subject condition averaged RMS-normalized response amplitudes were entered into a linear mixed-effects ANOVA model to evaluate how SSRP-induced changes in VEP amplitude were influenced by attentional deployment, session day, hemifield, and contrast level.

A single EEG session contributed 24 data points per participant—six contrast levels recorded before and six after SSRP induction for each hemifield. The model included fixed effects of pre/post (PrePost), contrast level, hemifield, attentional deployment and session day along with their n-way interactions, and accounted for repeated measures within participants by modeling subject-level variability as a random effect. This approach allowed us to examine multiple experimental factors simultaneously while controlling for inter-individual variability and within-subject dependencies inherent in repeated-measures designs. All statistical analyses were conducted in R (Version 4.4.1; R Core Team, 2023), and the model was fitted using the lme4 package (Version 3.1.164; (Bates et al., 2015)). Fixed effects included pre/post responses, Attentional Deployment Condition, Hemifield, and Contrast Level. A random intercept for subject was included to account for inter-individual variability across 48 observations per participant.

## Results

In brief, ssVEP contrast response functions (CRFs) were measured in the potentiated (left) and non-potentiated control (right) hemifields before and after a 10-minute presentation of high-contrast checkerboard stimuli restricted to the potentiated hemifield to induce stimulus-specific response potentiation (SSRP) (**Fig. 1A**). During SSRP induction, participants allocated endogenous covert attention either towards or away from the potentiating stimulus (**Fig. 1B**). Participants completed the two attention conditions on separate days, the order of which was counterbalanced across participants. All reported statistics were derived from a linear mixed-effects ANOVA model that included pre- and post-induction responses, contrast level, hemifield, attentional deployment condition, and session day as fixed effects, with participant modeled as a random intercept.

### Interaction Between Attention and Session Day

Interestingly, the mixed-effects models ANOVA detected a significant three-way interaction between PrePost, Attentional Deployment, and SessionDay (F = 8.79, *p* = 0.003, n = 26 participants). This finding suggests that the sequence of attentional conditions (Day 1 Congruent, Day 2 Incongruent versus Day 1 Incongruent, Day 2 Congruent) influenced response potentiation. To visualize this interaction, data were separated by counterbalanced groups, and pre/post-SSRP CRFs were plotted by attentional condition and SessionDay for each hemifield and attention condition (**Fig. 2**). **Figure 2A** depicts data from **Group A**, participants who completed the Attention-Congruent condition during SessionDay 1 and the Attention-Incongruent condition during SessionDay 2 (n = 13). **Figure 2B** shows **Group B**, who completed the sessions in the opposite order (n = 13). On SessionDay 1, response amplitude increases post-SSRP were observed when attention was allocated to the high-contrast SSRP-induction stimulus (**Fig. 2A**, panels 1 and 3), and this occurred in both hemifields. When attention was allocated *away* from the induction stimulus, post-SSRP responses showed no potentiation (**Fig. 2B**, panels 1 and 3), and responses in the non-potentiated hemifield showed a *decreased* response post-SSRP (**Fig. 2B**, panel 3), suggesting a reduction in response driven by attending away from the SSRP-inducing stimulus. On SessionDay 2, post-SSRP response potentiation emerged in both groups, in both hemifields, indicating that potentiation became more consistent across conditions on the second session (**Fig. 3**, cols. 2 and 4). This trend can be seen in individual participant data (**Supp. Fig. 2**).

**Figure 2.**
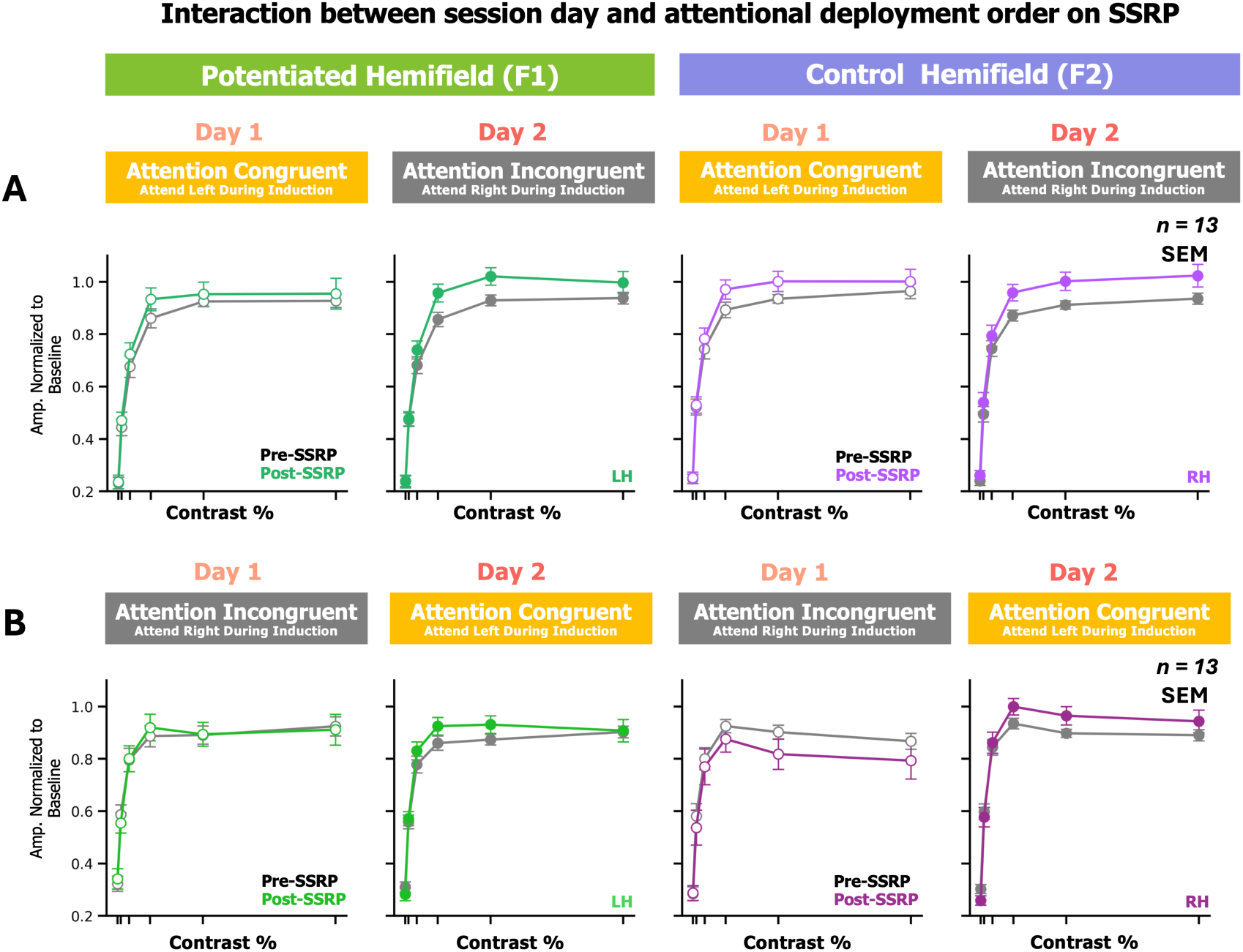
| Attention and Session Day independently enhance stimulus-specific response potentiation. Participants were separated into two counterbalanced groups based on the order of attentional deployment conditions completed across session days. The y-axis represents amplitude normalized to baseline, and the x-axis denotes contrast level. **Group A** (n =13, top row) completed the Congruent condition (attending toward the induction stimulus during SSRP) for the 1st session and Incongruent (attending away from the induction stimulus) for the 2nd session. **Group B** (n =13, bottom row) completed the conditions in the reverse order (Incongruent first, Congruent second). SessionDay 1 is denoted by white filled dots along the CRF (cols. 1 and 3), while SessionDay 2 is represented by filled dots (cols. 2 and 4). Hemifield is depicted in green for potentiated (F1) (cols. 1 and 2) and purple for non-potentiated (F2) (cols. 3 and 4). Attentional deployment conditions are denoted by the yellow and gray headers. On SessionDay 1 (cols. 1 and 3), response amplitudes increased in the Congruent condition (top row, panels 1 and 3) but not the Incongruent condition (bottom row, panels 1 and 3), with a trend toward a paradoxical decrease in the response in the non-potentiated hemifield in this condition. On SessionDay 2 (cols. 2 and 4), response amplitudes increased regardless of the attention condition in both hemifields.

To evaluate the differential effects by day, linear mixed effects models ANOVAs were performed on split SessionDay 1 and SessionDay 2 data. This confirmed a significant attentional modulation of SSRP on SessionDay 1 (PrePost x Attention: F = 9, *p* = 0.002) that was not present on SessionDay 2 (PrePost x Attention: *p* = 0.17). Attentional effects on SSRP were not hemifield-specific in split SessionDay 1 data (PrePost x Attention x Hemifield: *p* = 0.15). Our results conditionally confirm both **Hypothesis II and III,** indicating that attention enhances the expression of SSRP in naive observers, but accumulated effects across days predominate and occlude attentional effects in subsequent days of plasticity induction. Other ANOVA results will be discussed in the following sections.

### Effect of attention

Beyond PrePost x SessionDay x Attention interaction described above, other PrePost x Attention interactions were non-significant (PrePost × Attention × Hemifield: *p* = 0.46; PrePost × Attention: F = 0.98, *p* = 0.32). This confirmed that the attentional effect on SSRP is contextualized to naive observers and is not hemifield-specific.

### Accumulation across days

The effect of SessionDay was more pronounced and by itself statistically significant (PrePost x SessionDay: F = 4.45, *p* = 0.035). No significant three-way interactions were observed with Hemifield (PrePost × SessionDay × Hemifield: *p* = 0.46) or Contrast Level (PrePost × SessionDay × Contrast Level: *p* = 0.52), indicating that the enhanced potentiation effect on SessionDay 2 was neither hemifield-nor contrast-specific.

The increase in SSRP on SessionDay 2 could be due to either a decrease in the pre-SSRP response or an increase in the post-SSRP response, as in our initial analysis responses were normalized to the pre-SSRP response maximum on each day (see Methods). To distinguish these possibilities, we re-normalized each participant’s CRFs across both sessions to their SessionDay 1 pre-SSRP max response amplitude. **Supp. Fig. 3** shows that increased SSRP on SessionDay 2 reflected a combination of a slight decrease in pre-SSRP amplitude and a modest increase in post-SSRP amplitude relative to SessionDay 1 (cols. 1 and 3). We also examined whether the interval between sessions influenced the magnitude of SSRP accumulation. Although there was a weak trend toward greater potentiation on SessionDay 2 when sessions were spaced more than four days apart (**Supp. Fig. 4**), an ANOVA comparing pre-to post-SSRP response changes across interval groups revealed no significant effect.

### Retinotopic and contrast specificity

In addition to the lack of hemifield and contrast specificity in the above interactions, VEP potentiation was neither contrast-specific (Pre-Post × Contrast: *p* = 0.16) nor hemifield-specific (PrePost × Hemifield: *p* = 0.56), weighing against Hypothesis I that SSRP demonstrates LTP-like input specificity.

### First-level ANOVA factors

As expected, the ANOVA revealed a significant overall effect of SSRP, with response amplitudes increasing after SSRP compared to baseline (F =10, *p* = 0.002) (**Supp. Fig. 1B**). Also as expected, there was a significant primary effect of Contrast (F = 845, *p* < 0.001).

## Discussion

The primary finding of this study was a significant three-way interaction between PrePost, Attentional Deployment, and SessionDay, indicating that attention enhances SSRP in naive observers (Hypothesis II confirmed), but across-day accumulation of SSRP effects predominates in experienced observers (Hypothesis III confirmed) and occludes any attentional modulation of visual cortical plasticity. During the initial exposure, directing attention toward the potentiating stimulus facilitated SSRP, whereas attending away from the potentiating stimulus caused a paradoxical reduction of response in the control hemifield. The enhancement of post-SSRP responses across both attentional conditions and hemifields on the second day indicates that repeated exposure played the dominant role in facilitating potentiation across sessions. Finally, contrary to some prior reports and our Hypothesis I, in our paradigm SSRP was not retinotopic-or contrast-specific. Although animal studies of SSRP have shown exceptional input specificity as reflected by spatial frequency and orientation selectivity (Cooke and Bear, 2010, Kaneko et al., 2017), and a related form of SSRP called tetanic stimulation does show reports of retinotopic and orientation specificity in humans (Teyler et al., 2005, McNair et al., 2006), Normann et al.’s (2007) paradigm which elicits SSRP in humans, has not been systematically tested for input specificity. In our hemifield-specific design, we found no evidence of input specificity, as SSRP occurred in both hemifields despite induction being restricted to the left hemifield. To our knowledge, this is the first study to evaluate SSRP effects in humans using a repeated-exposure paradigm analogous to animal models of SSRP. Although response potentiation appeared somewhat stronger at intermediate-to-high contrasts, this trend did not reach statistical significance, indicating no reliable contrast-dependent input specificity under the present protocol. In the future, a model-based analysis of contrast-response function shape may allow determination if SSRP enhances activity via contrast-gain- or response-gain-like mechanisms, an analysis that could allude to differential plasticity in inhibitory and excitatory visual cortical circuits (Ling and Carrasco, 2006). Our results tentatively suggest that stimulus-specific response potentiation may be a misnomer in the case of visual cortical plasticity induced by 10 minutes of repetitive sensory stimulation in humans. Future work employing spatial-frequency or orientation sweeps could more directly probe feature-specific channels and permit finer comparisons with rodent SSRP studies.

Plasticity processes are known to unfold gradually across sessions or days, as shown in spaced LTP paradigms (Frenkel et al., 2006, Cooke and Bear, 2010), and multi-day perceptual learning studies reporting large gains in perceptual sensitivity with repeated training (Furmanski et al., 2004, Cooke and Bear, 2010, Hua et al., 2010, Kaneko et al., 2017). Consistent with this literature, our findings suggest that SSRP may increase across days in human participants. There is a rough similarity between our findings of enhanced SSRP on day 2, to the accumulation of response amplitude enhancements observed across multiple days of SSRP effects in rodents (Frenkel et al., 2006). However, the in-rodent studies did not split their measures into a pre- and post-SSRP analyses, rather pooling all responses per day to show a net increase in response amplitude across days. Our data, in contrast, show a larger within-day change on the second day of recording, and we do not see an overall increase in SessionDay 2 responses when normalizing to SessionDay 1 (**Supp. Fig. 3**). The precise relationship between the across-day plasticity effects we report here and previous in-rodent studies will require clarification in future studies. While attention modestly facilitated SSRP expression on SessionDay 1, its influence diminished by SessionDay 2 despite stronger overall potentiation, highlighting a dissociation between transient attentional modulation and sustained, experience-driven plasticity. This pattern suggests that attention may enhance early stages of plasticity, particularly in naïve circuits, but that repeated stimulation can elicit lasting changes independently. Rodent studies have posited a potential cholinergic contribution to attention-driven modulation of SSRP (Kaneko et al., 2017), which could be assessed in our paradigm with co-administration of acetylcholine modulators like donepezil (Rokem et al., 2010), an interesting avenue for further research.

One unexpected observation that deserves further exploration is the decrease in response amplitude within non-potentiated hemifield on SessionDay 1 in the Incongruent condition (See **Fig. 2B**, panel 3). This effect could reflect a non-local influence of the induction stimulus or, more intriguingly, a long-term depression (LTD)-like process driven by the low-contrast (2%) control stimulus that is modulated by attention. The consensus view in the field is that a high-contrast stimulus is necessary to induce SSRP, although we are not aware of a study that assessed VEP plasticity effects of repeated presentations of low contrast stimuli. A possibly analogous attentional modulation of LTD-like plasticity has been observed in the motor cortex (Kamke et al., 2012). Future experiments can be designed to distinguish these different possibilities.

The paradigm used in the current study was primarily inspired by the protocol originally described by Normann et al. (Normann et al., 2007) and shown to have a robust effect size in the large-scale study conducted by the Elvsåshagen group (Valstad et al., 2020). For the sake of comparison, the distinctions between the two protocols will be delineated here. In the Normann et al. paradigm, the pre-post-SSRP stimulus consisted of a full-field 100% contrast checkerboard stimulus, 0.5° spatial frequency, sign reversing at 2 repetitions per cycle (rps). The current study’s pre-post-SSRP stimulus differed in that checkerboards (approximately 0.5° SF) were semicircular extending to 10° with checks eccentricity scaled; stimuli were presented at 6 and 7.5 Hz; and contrast was swept from 2 to 100% contrast. The induction stimulus of Norman et al. (Normann et al., 2007) was the same stimulus as pre/post but presented continuously for ten minutes. The current study’s induction stimulus was similar to our pre/post stimulus but with a sign reversal rate and contrast matching Norman et al (2 Hz and 100%), but differed in that it was only presented in the left hemifield. In Normann et al., there was a single baseline 20-second trial, and 20-second post-trials were presented at 2, 8, 12, 18, 22, and 28 minutes after the SSRP block. In the current study, in contrast, thirty-nine 10.66-second trials were presented pre- and post-, and post-stimuli were presented starting 3-4 minutes after the end of the SSRP block. We note that our paradigm may have missed delayed phases of potentiation that emerge tens of minutes later (Valstad et al., 2020).

### Limitations

While this study advances our understanding of stimulus-specific response potentiation (SSRP), it also highlights several limitations that inform future work: 1) Our study did not include eye-tracking, so we are not able to precisely verify fixation in the induction block. Weighing against the possibility of this impacting our results, a post-hoc analysis of the electrooculogram recordings indicated that horizontal eye movements during the induction block were minimal. Moreover, our primary observed effect of enhanced response potentiation on Session-Day 2 would not be affected by issues with fixation. 2) Participants may not have been strongly allocating attention in the task. We think this is not a major contributor to our results, because we assessed task response consistency as a rough corollary of attention, and when we excluded participants who were not performing the task consistently, it did not alter the results. However, in future studies, a more rigorously designed attentional task with better behavioral measures and eye tracking may be more sensitive at detecting attentional effects on SSRP. 3) The behavioral tasks implemented used during the pre-/post-blocks were primarily designed to enforce fixation rather than probe SSRP-related changes, which limits their usefulness in capturing behaviorally relevant effects of SSRP. In future studies a contrast discrimination task at threshold could be performed by participants pre-post-SSRP to compare electrophysiological plasticity effects to perceptual plasticity effects. 4) Participants were not actually allocating attention to the potentiating stimulus. Instead they were allocating attention to the retinotopic area that encompassed the SSRP stimulus, but were trying to detect small patches of subtle contrast change within that retinotopic extent. This paradigm is known to enhance ssVEP responses to the background stimulus. However, from this one perspective, the potentiating stimulus may serve as a distractor in relation to the attended targets. Therefore the attentional system may have been trying to suppress or filter out responses to the potentiating stimulus in our paradigm, which could paradoxically suppress SSRP effects. This interesting possibility will be assessed as well in future experiments. 5) The low-contrast stimulus presented in the control right hemifield during induction may have elicited its own plasticity effects (as described above). If the low-contrast control stimulus in our paradigm did induce some form of plasticity, perhaps this is evident in the attention-dependent SessionDay 1 response depression we observed. Future studies can assess this interesting possibility.

Finally, we can provide no comprehensive mechanistic theory to explain how variables such as stimulus contrast, repetitions per cycle, stimulus type, and eccentricity degree facilitate SSRP susceptibility. The design of our SSRP experiment remains largely empirical, with no theoretical framework to account for how temporal frequency or duration of SSRP blocks impacts the magnitude or persistence of response enhancement. This gap in mechanistic understanding highlights the critical need for further investigation into the visual system’s plasticity mechanisms and their interactions with specific stimulus properties overall.

## Supplementary Materials

**Supplemental Figure 1.**
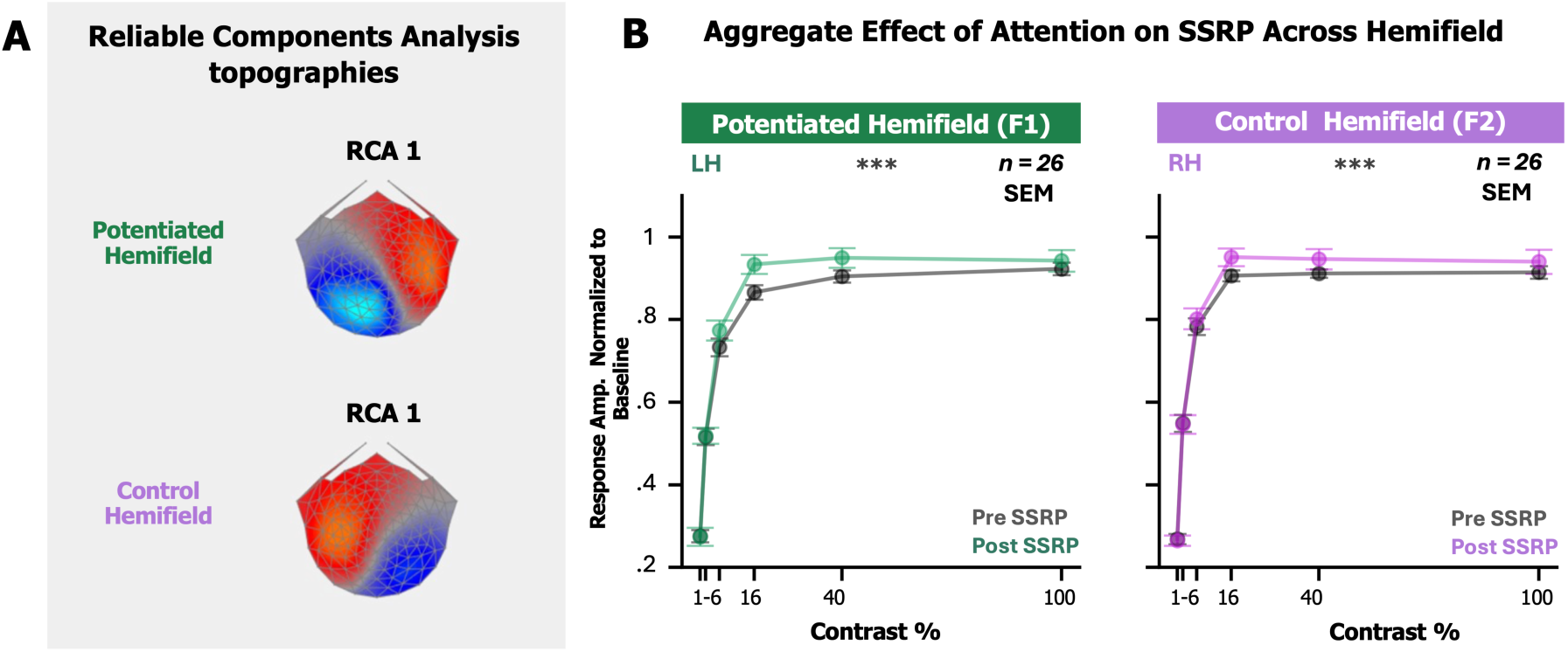
| RCA topography and average Pre- Post- CRF per hemifield. **A** Top-view topological maps (nose facing up) show the electrode weights obtained from the first RCA component for each hemifield stimulus. Notably, for each hemifield frequency tag, the first RCA component falsely localizes to the ipsilateral occipital electrodes, presumably due to dipole cancellation between the inferior and superior occipital sulci. **B** Mean contrast response functions before (black) and after (colored) the induction block in the potentiated (green) and control (purple) hemifields, data pooled across attentional conditions and session days Error bars depict S.E.M., n = 26 participants. The y-axis indicates response amplitude normalized per-participant to pre-SSRP response-max, and the x-axis represents visual contrast. ** F = 10, *p* = 0.002, Linear mixed-effects-models ANOVA Pre vs. Post SSRP.

**Supplemental Figure 2.**
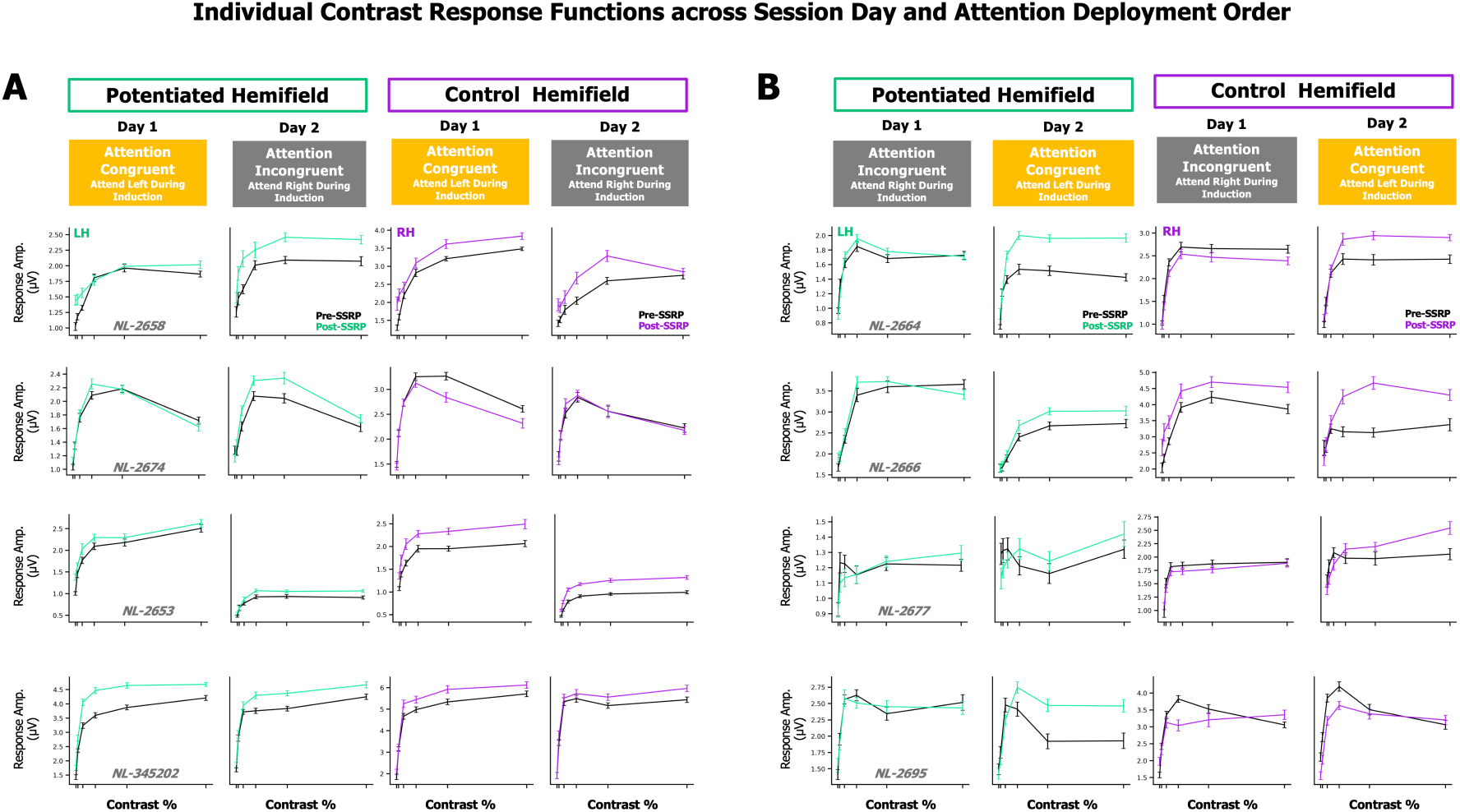
| Individual CRFs across session day and attention deployment order. Each row represents individual RMS Contrast Response Functions (mean, S.E) measured Pre-SSRP (black) and Post-SSRP (colored) development for the potentiated (F1) and control (F2) hemifields across SessionDay 1, SessionDay 2 and the Attentional Deployment condition completed for the respective SessionDay. The y-axis represents amplitude, and the x-axis denotes contrast level. Individual data are divided into two groups based on the counterbalanced order of attentional deployment conditions: **Group A** completed the **Attention SSRP Congruent condition for the 1st session**, followed by completing the **Attention SSRP Incongruent condition for the 2nd session**. **Group B** completed these conditions in the reverse order. Overall between both groups, potentiation Post-SSRP on the second is much more pronounced compared to first session responses for both hemifields regardless of attentional deployment order, especially at higher contrasts (cols. 2 and 4).

**Supplemental Figure 3.**
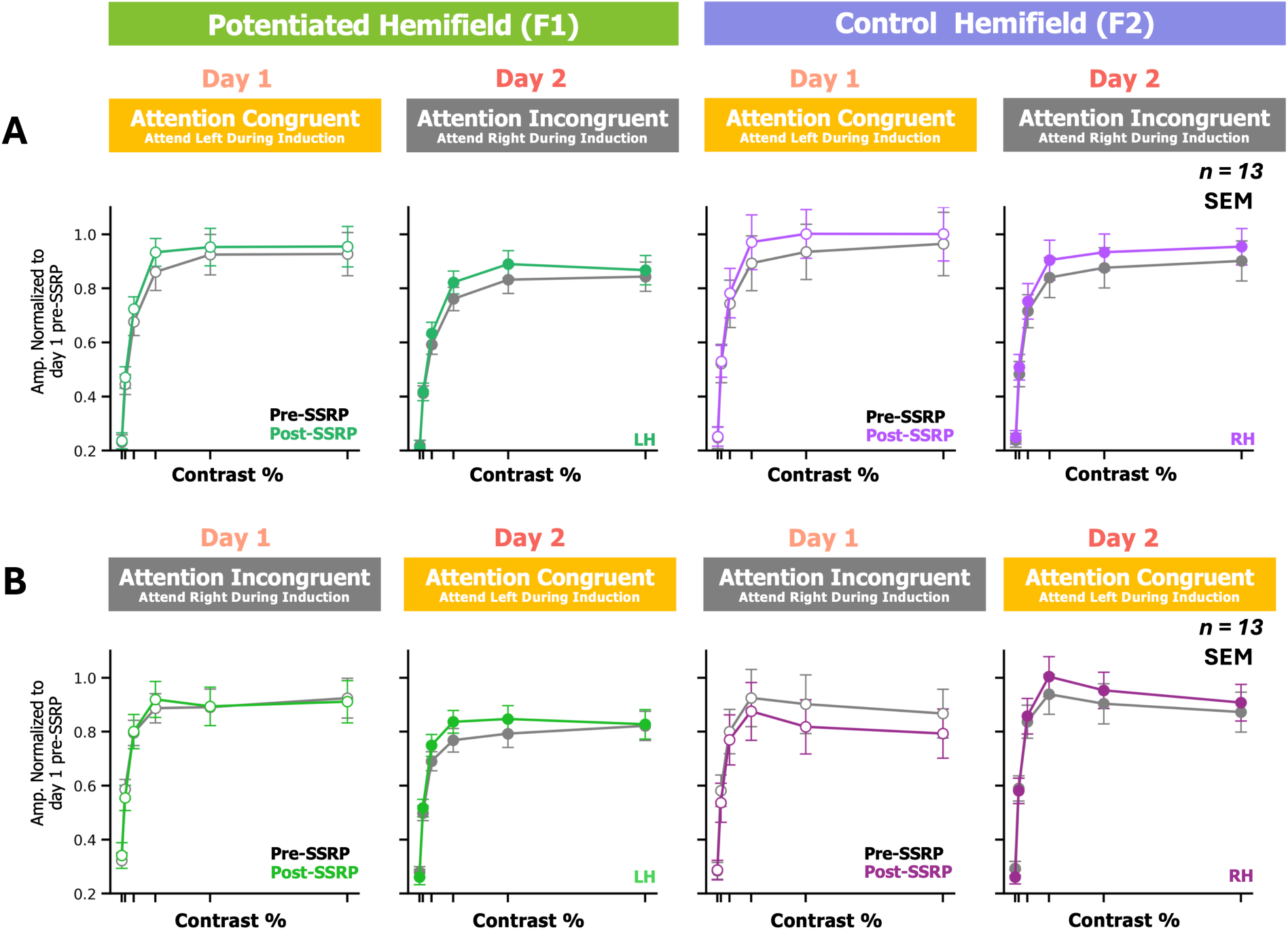
| Data from Figure 2, normalized to Day 1 Pre baseline responses to visualize across day changes in response amplitude. Plotting the data in this fashion reveals that enhanced responses on SessionDay2 were due to a combination of reduced pre-SSRP responses and augmented post-SSRP responses (cols. 2 and 4). This suggests that some level of adaptation and plasticity affected CRF development across the first and second session day. After running the same LME model on this version of the normalized data, the PrePost × SessionDay × AttentionDeployment interaction did not reach significance.

**Supplemental Figure 4.**
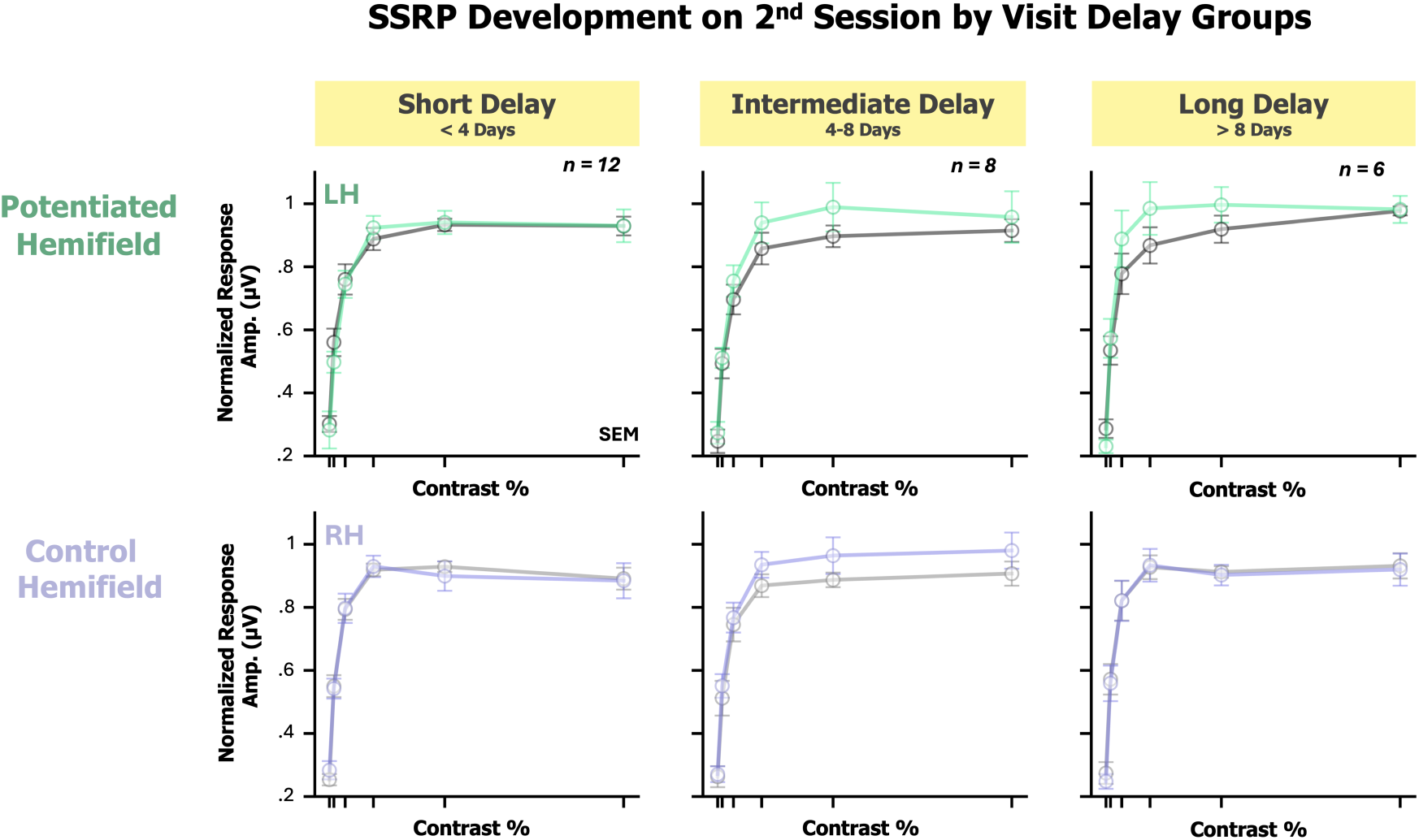
| Session Day 2 contrast-response functions as function of delay time. The y-axis represents amplitude normalized baseline, and the x-axis denotes contrast level. Pre-SSRP (black) and post-SSRP, (green/purple) CRF’s (mean, S.E.M) are shown based on different session delay visit groups: Short Delay (2nd visit was less than 4 days, 1st column), Intermediate Delay (2nd visit was 4-8 days, 2nd column), Long Delay (2nd visit was more than 8 days later, 3rd column). The first row shows responses Pre-Post SSRP in the potentiated hemifield, regardless of attentional deployment condition. The second row shows responses Pre-Post SSRP in the control hemifield in the respective day delay group. The potentiated hemifield seems to show stronger mid-range contrast enhancement (row 1), but this was not significant within or across groups, so enhancement did not increase with time. Still, the sample size was small, and delay between sessions was not systematically controlled.

## References

Justin M. Ales, Jacob L. Yates, and Anthony M. Norcia. V1 is not uniquely identified by polarity reversals of responses to upper and lower visual field stimuli. NeuroImage, 52(4):1401–1409, 5 2010. 10.1016/j.neuroimage.2010.05.016. URL https://doi.org/10.1016/j.neuroimage.2010.05.016.

Ryan T. Ash, Kerry C. Nix, and Anthony M. Norcia. Stability of steady-state visual evoked potential contrast response functions. Psychophysiology, 61(1), 8 2023. 10.1111/psyp.14412. URL https://doi.org/10.1111/psyp.14412.

Douglas Bates, Martin Mächler, Ben Bolker, and Steve Walker. Fitting linear mixed-effects models using lme4. Journal of Statistical Software, 67(1):1–48, 2015. 10.18637/jss.v067.i01. URL https://doi.org/10.18637/jss.v067.i01.

Marisa Carrasco. Visual attention: The past 25 years. Vision Research, 51(13):14841525, jul 2011. 10.1016/j.visres.2011.04.012. URL https://doi.org/10.1016/J.visres.2011.04.012.

Matthew R. Cavanaugh, Duje Tadin, Marisa Carrasco, and Krystel R. Huxlin. Benefits of endogenous spatial attention during visual double-training in cortically-blinded fields. Frontiers in Neuroscience, 16:771623, 2022. ISSN 1662-453X. 10.3389/fnins.2022.771623. URL https://doi.org/10.3389/fnins.2022.771623.

Sam F. Cooke and Mark F. Bear. Visual experience induces long-term potentiation in the primary visual cortex. The Journal of Neuroscience, 30(48):16304–16313, December 2010. 10.1523/JNEUROSCI.4333-10.2010.

Sam F. Cooke and Mark F. Bear. Stimulus-Selective Response Plasticity in the Visual Cortex: An Assay for the Assessment of Pathophysiology and Treatment of Cognitive Impairment Associated with Psychiatric Disorders. Biological Psychiatry, 71(6):487–495, 10 2011. 10.1016/j.biopsych.2011.09.006. URL https://doi.org/10.1016/j.biopsych.2011.09.006.

J. W. Dias, C. M. McClaskey, J. A. Rumschlag, and K. C. Harris. Sensory tetanisation to induce long-term-potentiation-like plasticity: A review and reassessment of the approach. European Journal of Neuroscience, Volume 56(12) – 2022: 6115–6140, 2022. ISSN 0953-816X. 10.1111/ejn.15847. URL https://doi.org/10.1111/ejn.15847.

Jacek P Dmochowski, Alex S Greaves, and Anthony M Norcia. Maximally reliable spatial filtering of steady state visual evoked potentials. NeuroImage, 109:63–72, 1 2015. 10.1016/j.neuroimage.2014.12.078. URL https://doi.org/10.1016/J.neuroimage.2014.12.078.

Ian Donovan and Marisa Carrasco. Endogenous spatial attention during perceptual learning facilitates location transfer. Journal of Vision, 18(11):7, 2018. 10.1167/18.11.7. URL https://doi.org/10.1167/18.11.7.

Ian Donovan, Angela Shen, Cristina Tortarolo, Antoine Barbot, and Marisa Carrasco. Exogenous attention facilitates perceptual learning in visual acuity to untrained stimulus locations and features. Journal of Vision, 20(4):18–18, 04 2020. ISSN 1534-7362. 10.1167/jov.20.4.18. URL https://doi.org/10.1167/Jov.20.4.18.

Mikhail Y. Frenkel, Nathaniel B. Sawtell, Antonia Cinira M. Diogo, Bongjune Yoon, Rachael L. Neve, and Mark F. Bear. Instructive effect of visual experience in mouse visual cortex. Neuron, 51(3):339–349, 8 2006. 10.1016/j.neuron.2006.06.026. URL https://doi.org/10.1016/j.neuron.2006.06.026.

Christopher S. Furmanski, Denis Schluppeck, and Stephen A. Engel. Learning strengthens the response of primary visual cortex to simple patterns. Current Biology, 14(7):573–578, April 2004. 10.1016/j.cub.2004.03.032.

Tianmiao Hua, Pinglei Bao, Chang-Bing Huang, Zhenhua Wang, Jinwang Xu, Yifeng Zhou, and Zhong-Lin Lu. Perceptual learning improves contrast sensitivity of v1 neurons in cats. Current Biology, 20(10):887–894, May 2010. 10.1016/j.cub.2010.03.066.

Sharon C. Hung and Marisa Carrasco. Feature-based attention enables robust, long-lasting location transfer in human perceptual learning. Scientific Reports, 11(1): 13914, July 2021. 10.1038/s41598-021-93016-y. Erratum: Scientific Reports. 2021 Aug 23;11(1):17293. doi:10.1038/s41598-021-96732-7.

Michael S. Jacob, Brian J. Roach, Holly K. Hamilton, Ricardo E. Carrión, Aysenil Belger, Erica Duncan, Jason Johannesen, Matcheri Keshavan, Sandra Loo, Margaret Niznikiewicz, Jean Addington, Carrie E. Bearden, Kristin S. Cadenhead, Tyrone D. Cannon, Barbara A. Cornblatt, Thomas H. McGlashan, Diana O. Perkins, William Stone, Ming Tsuang, Elaine F. Walker, Scott W. Woods, and Daniel H. Mathalon. Visual cortical plasticity and the risk for psychosis: An interim analysis of the North American Prodrome Longitudinal Study. Schizophrenia Research, 230:26–37, 3 2021b. 10.1016/j.schres.2021.01.028. URL https://doi.org/10.1016/j.schres.2021.01.028.

Marc R. Kamke, Michelle G. Hall, Hayley F. Lye, Martin V. Sale, Laura R. Fenlon, Timothy J. Carroll, Stephan Riek, and Jason B. Mattingley. Visual attentional load influences plasticity in the human motor cortex. Journal of Neuroscience, 32 (20):7001–7008, 5 2012. 10.1523/jneurosci.1028-12.2012. URL https://doi.org/10.1523/jneurosci.1028-12.2012.

Marc R. Kamke, Alexander E. Ryan, Martin V. Sale, Megan E. J. Campbell, Stephan Riek, Timothy J. Carroll, and Jason B. Mattingley. Visual spatial attention has opposite effects on bidirectional plasticity in the human motor cortex. Journal of Neuroscience, 34(4):1475–1480, 1 2014. 10.1523/jneurosci.1595-13.2014. URL https://doi.org/10.1523/jneurosci.1595-13.2014.

Megumi Kaneko, Yu Fu, and Michael P. Stryker. Locomotion induces stimulus-specific response enhancement in adult visual cortex. The Journal of Neuroscience, 37(13):3532–3543, March 2017. 10.1523/JNEUROSCI.3760-16.2017.

Ian J. Kirk, Nicholas A. McNair, John P. Hamm, Wesley C. Clapp, Daniel H. Mathalon, Ihsan Cavus, and Timothy J. Teyler. Long-term potentiation (ltp) of human visual evoked responses. Brain Research Bulletin, 76(5):509–516, 2008. ISSN 0361-9230. 10.1016/j.brainresbull.2008.01.021. URL https://doi.org/10.1016/j.brainresbull.2008.01.021.

Ian J. Kirk, Meg J. Spriggs, and Rachael L. Sumner. Human eeg and the mechanisms of memory: investigating long-term potentiation (ltp) in sensory-evoked potentials. Journal of the Royal Society of New Zealand, 51(1):24–40, 2021. 10.1080/03036758.2020.1780274. URL https://doi.org/10.1080/03036758.2020.1780274.

Stefan Klöppel, Eliza Lauer, Jessica Peter, Lora Minkova, Christoph Nissen, Claus Normann, Janine Reis, Florian Mainberger, Michael Bach, and Jacob Lahr. Ltp-like plasticity in the visual system and in the motor system appear related in young and healthy subjects. Frontiers in Human Neuroscience, 9:506, 2015. 10.3389/fnhum.2015.00506. URL https://doi.org/10.3389/fnhum.2015.00506.

Sam Ling and Marisa Carrasco. Sustained and transient covert attention enhance the signal via different contrast response functions. Vision Research, 46(8-9): 1210–1220, 2006. 10.1016/j.visres.2005.05.008.

Zhong-Lin Lu, Tianmiao Hua, Chang-Bing Huang, Yifeng Zhou, and Barbara Anne Dosher. Visual perceptual learning. Neurobiology of Learning and Memory, 95(2):145–151, 2011. ISSN 1074-7427. 10.1016/j.nlm.2010.09.010. URL https://www.sciencedirect.com/science/article/pii/ S1074742710001644. RF Thompson Special Issue.

Nicolas A. McNair, Wesley C. Clapp, Jeff P. Hamm, Timothy J. Teyler, Michael C. Corballis, and Ian J. Kirk. Spatial frequency-specific potentiation of human visual-evoked potentials. NeuroReport, 17(7):739–741, May 2006. 10.1097/01.wnr.0000215775.53732.9f.

Steven T. Morgan, J. Chris Hansen, and Steven A. Hillyard. Selective attention to stimulus location modulates the steady-state visual evoked potential. Proceedings of the National Academy of Sciences of the United States of America, 93(10):4770– 4774, 1996. ISSN 0027-8424. 10.1073/pnas.93.10.4770. URL https://doi.org/10.1073/pnas.93.10.4770.

Bahadur K. Kesavabhotla K. Ungerleider L. G. Mukai, I. Exogenous and endogenous attention during perceptual learning differentially affect post-training target thresholds. Journal of Vision, 11(1):25, 2011. 10.1167/11.1.25. URL https://doi.org/10.1167/11.1.25.

M. M. Müller, T. W. Picton, P. Valdés-Sosa, J. Riera, W. A. Teder-Sälejärvi, and S. A. Hillyard. Effects of spatial selective attention on the steady-state visual evoked potential in the 20–28 hz range. Brain Research: Cognitive Brain Research, 6(4):249–261, 1998. 10.1016/S0926-6410(97)00036-0. URL https://doi.org/10.1016/S0926-6410(97)00036-0.

Kieu Ngoc Nguyen, Takeo Watanabe, and George John Andersen. Role of en-dogenous and exogenous attention in task-relevant visual perceptual learning. PLOS ONE, 15(8):e0237912, aug 2020. 10.1371/journal.pone.0237912. URL https://doi.org/10.1371/journal.pone.0237912.

Anthony M. Norcia, L. Gregory Appelbaum, Justin M. Ales, Benoît R. Cottereau, and Bruno Rossion. The steady-state visual evoked potential in vision research: A review. Journal of Vision, 15(6):4, 2015. 10.1167/15.6.4. URL https://doi.org/10.1167/15.6.4.

Claus Normann, Daniela Schmitz, Andrea Fürmaier, Carsten Döing, and Michael Bach. Long-Term Plasticity of Visually Evoked Potentials in Humans is Altered in Major Depression. Biological Psychiatry, 62(5):373–380, 1 2007. 10.1016/j.biopsych.2006.10.006. URL https://doi.org/10.1016/j.biopsych.2006.10.006.

Ariel Rokem, Ayelet N. Landau, Dave Garg, William Prinzmetal, and Michael A. Silver. Cholinergic enhancement increases the effects of voluntary attention but does not affect involuntary attention. Neuropsychopharmacology, 35(13):2538– 2544, 2010. 10.1038/npp.2010.118.

Katja Stefan, Matthias Wycislo, and Joseph Classen. Modulation of associative human motor cortical plasticity by attention. Journal of Neurophysiology, 92 (1):66–72, 6 2004. 10.1152/jn.00383.2003. URL https://doi.org/10.1152/jn.00383.2003.

Rachel L. Sumner, Matthew J. Spriggs, Suresh D. Muthukumaraswamy, and Ian J. Kirk. The role of hebbian learning in human perception: a methodological and theoretical review of the human visual long-term potentiation paradigm. Neuroscience & Biobehavioral Reviews, Volume 115–2020, 2020. ISSN 0149-7634. 10.1016/j.neubiorev.2020.03.013. URL https://doi.org/10.1016/j.neubiorev.2020.03.013.

Sarit Felicia Anais Szpiro and Marisa Carrasco. Exogenous attention enables perceptual learning. Psychological Science, 26:1854 –1862, 2015. URL https://api.semanticscholar.org/CorpusID:13000477.

Timothy J. Teyler, Jeff P. Hamm, Wesley C. Clapp, Blake W. Johnson, Michael C. Corballis, and Ian J. Kirk. Long-term potentiation of human visual evoked responses. The European Journal of Neuroscience, 21(7):2045–2050, April 2005. 10.1111/j.1460-9568.2005.04007.x.

Mathias Valstad, Torgeir Moberget, Daniël Roelfs, Nora B. Slapø, Clara M.F. Timpe, Dani Beck, Geneviève Richard, Linn Sofie Sæther, Beathe Haatveit, Knut Andre Skaug, Jan Egil Nordvik, Christoffer Hatlestad-Hall, Gaute T. Einevoll, Tuomo Mäki-Marttunen, Lars T. Westlye, Erik G. Jönsson, Ole A. An-dreassen, and Torbjørn Elvsåshagen. Experience-dependent modulation of the visual evoked potential: Testing effect sizes, retention over time, and associations with age in 415 healthy individuals. NeuroImage, 223:117302, 2020. ISSN 1053-8119. 10.1016/j.neuroimage.2020.117302. URL https://www.sciencedirect.com/science/article/pii/S1053811920307886.

Mathias Valstad, Daniël Roelfs, Nora B Slapø, Clara M F Timpe, Ahsan Rai, Anna Maria Matziorinis, Dani Beck, Geneviève Richard, Linn Sofie Sæther, Beathe Haatveit, Jan Egil Nordvik, Christoffer Hatlestad-Hall, Gaute T Einevoll, Tuomo Mäki-Marttunen, Marit Haram, Torill Ueland, Trine V Lagerberg, Nils Eiel Steen, Ingrid Melle, Lars T Westlye, Erik G Jönsson, Ole A Andreassen, Torgeir Moberget, and Torbjørn Elvsåshagen. Evidence for Reduced Long-Term Potentiation-Like Visual Cortical plasticity in schizophrenia and bipolar disorder. Schizophrenia Bulletin, 47(6):1751–1760, 4 2021. 10.1093/schbul/sbab049. URL https://doi.org/10.1093/schbul/sbab049.

Idil Çavuş, Robert M. G. Reinhart, Brian J. Roach, Ralitza Gueorguieva, Timothy J. Teyler, Wesley C. Clapp, Judith M. Ford, John H. Krystal, and Daniel H. Math-alon. Impaired visual cortical plasticity in schizophrenia. Biological Psychiatry, 71(6):512–520, 2012. 10.1016/j.biopsych.2012.01.013.

